# Single-cell transcriptomics and mouse model phenotyping for biomarker screen of peripheral blood biomarkers in Huntington’s disease

**DOI:** 10.64898/2025.12.14.694089

**Authors:** Paula Martín-Climent, Samanta Ortuño-Miquel, Juan F. Gallego-Serna, Silvia Martí-Martínez, Luis M. Valor

## Abstract

Transcriptional dysregulation is among the most prominent molecular alterations in Huntington’s disease (HD). It is not confined to brain but is also extended to peripheral tissues and cells, enabling minimally invasive screens aimed at identifying transcriptional surrogates of the health status in HD mutation carriers. Nonetheless, transcriptomics approaches have failed to identify consistent candidates from peripheral blood, probably due to the low impact of the HD mutation in the transcriptional profiles of circulating cells that can be masked by the high cellular complexity of this biofluid. In this study, we applied for the first time single-cell RNA-seq in peripheral blood mononuclear cells (PBMCs) to determine those cells that accumulate the most prominent gene expression changes as potential sources of reliable biomarkers. We observed both specific and common transcriptional alterations across different blood cell subtypes, justifying partial overlapping of our results with published bulk transcriptomics datasets. To relate these gene expression patterns with disease progression in the absence of a large cohort of patients, we examined selected candidates in a phenotypically characterized cohort of transgenic R6/1 mice. Of the tested genes, only the changes in the interferon related gene *Irf7* in blood were correlated with motor coordination performance in mutant mice. Notably, this gene was significantly upregulated in the striatum of these animals. Overall, transcriptional-based changes in peripheral blood can be linked to HD although dissection of individual blood cells in combination with subsequent validation in mouse models emphasizes the challenges in obtaining clinically relevant biomarkers in HD.

## Introduction

Huntington’s disease (HD) is a highly disabling condition with lethal consequences that is caused by an aberrant expansion of CAG triplets in the Huntingtin (*HTT*) gene. While waiting for an effective therapy, HD mutation carriers are in need of specific biomarkers to report their real health status, to predict the onset and progression of the diverse manifestation of symptoms, and to evaluate the response to therapeutic regimes; however, our current knowledge of biomarkers is not being applied to HD clinical management [1]. Regarding the potential sources of biomarkers, peripheral blood obtained through venopunctions is among the most preferred due to its minimal invasibility and maximal feasibility and universality, with the potential to inform about both systemic and brain-derived pathological perturbations since neural subproducts can be released into the bloodstream.

We and others have extensively reviewed the relevance of transcriptional dysregulation in HD [2–4], an early phenomenon that is not restricted to the most affected brain regions by the *HTT* mutation (e.g., basal ganglia and associated cortical areas) but it can also be detected peripherally (e.g., liver, muscle, adipocyte tissue, blood, skin), further supporting the nonneuronal clinical symptomatology of HD (reviewed in [1]). Since the beginning of modern HD research, transcriptomics technologies have been applied to screen accessible and quantifiable variations of gene expression with the potential to be linked to different disease stages in the peripheral blood from mutation carriers, and to be anticipated to the onset of declared symptomatology in sufficient time to facilitate decision-making [1]. However, the retrieval of clinically relevant candidates has been proven to be difficult. Despite using validation cohorts within each study, the proposed candidates were at best partially confirmed in few independent reports, but without being consistent across studies. This is exemplified by efforts to summarize newly generated datasets at the time with previous results, in which overlapping genes were mainly detected in a pair-wise fashion [5,6]. These results indicated that transcriptional changes in HD blood were not sufficiently pronounced as in the brain to overcome the interindividual variability and demographic variations that are features of unsupervised populations in humans. This situation is exacerbated due to the usage of diverse transcriptomics platforms and analytical procedures that hinder direct comparisons across studies.

In this context, the high cellular heterogeneity of the blood may be masking the nuclear effects of *HTT* mutation, which can be critical if such changes are more prominent in certain cell subpopulations. Indeed, transcriptional dysregulation in blood is still fully justified as the expression of mutant HTT (mHTT) has been detected in both innate and adaptive immune cells [7] and circulating proinflammatory factors (e.g., IL-6, IL-8, TNF- α) are increased according to the disease stage [8–10]. To reach more definitive conclusions regarding transcriptional-based blood biomarkers in HD, we applied for the firs time single-cell transcriptomics (scRNA-seq) in peripheral blood from symptomatic volunteers to determine those disease-specific cell variants that are more susceptible to contain reliable biomarkers (i.e., which cell subpopulation(s) can accumulate most of the disease-dependent molecular alterations) and how biomarker candidates behave across different cell types, as isolation of specific subpopulations can augment the detection sensitivity (as reported for myeloid cells from patients and mouse models [11,12]). This knowledge can lead to further exploration of specific cellular targets using more affordable techniques in the clinics. Although the use of this costly technology is still not suitable for the study of large cohorts, scRNA-seq (or its single-nuclear variant snRNA-seq) is widening our understanding regarding HD pathogenesis by dissecting the defects occurring across multiple neuronal, glial and other non-neuronal cell types and illustrates the unforeseen cellular heterogeneity even in seemingly homogeneous subpopulations [13–16].

We complemented this approach by investigating the correlation between gene expression disruption and pathological phenotype in a mouse model, which was more feasible than the analysis of clinical data from large cohorts which are challenging to obtain for rare disorders, not normally supported by sample collections from initiatives such as Enroll-HD (https://www.enroll-hd.org) for biomarker research. Also, the use of animal models enables the comparison between brain and peripheral alterations, assuming that most of the alterations can be conserved across species (as it happens with transcriptional dysregulation in brain [17,18]). This work is a proof-of-concept study to assess the convenience of single cell transcriptomics for the proposal of novel candidates in biomarker screens and the use of phenotypically characterized animal models for their validation.

## Materials and Methods

### Human samples

The study was conducted according to the guidelines of the Declaration of Helsinki, following the national and regional laws and regulations concerning biomedical research on human samples, personal data protection and the use of biobank services, after approval by the local ethics committee: Comité de Ética de Investigación Clínica con medicamentos (CEIm) del Hospital General Universitario Dr. Balmis (HGUDrB), reference number PI2021/058, 12^th^ November 2021. Samples and clinical data were provided by the Biobank of ISABIAL which is adhered to the Spanish National Biobanks Network and integrated in the Valencian Biobanking Network, following standard operating procedures.

Human volunteers were recruited through the Service of Neurology of the HGUDrB, and consisted on 4 controls (without the HD mutation) and 4 HD symptomatic patients, with equal representation of both sexes in each condition (2 women and 2 men) and balanced age (median (IQR) = 52.0 (9.8) in controls, median (IQR) = 57.5 (8.8) in patients). The number of CAG repeats in the expanded allele was 43.5 (3.5) in the HD group. The stage of HD according to the Total Functional Capacity (TFC) [19] was 1 for all controls and 4-5 for all symptomatic patients.

Human blood samples were collected in EDTA-K2 Vaccutainer tubes (BD) for isolation of peripheral blood mononuclear cells (PBMC) after a density gradient centrifugation using Pancoll human,1.077 g/mL (PAN-Biotech) following this procedure: 1,300xg 15min, followed by a wash with 0.1M PBS 1,000xg 5min. Cells were immediately cryopreserved in two aliquots per donor using the Demonstrated Protocol CG00039 RevD “Fresh Frozen Human PBMC for scRNA-seq” (10xGenomics) that was previously validated in our laboratory to preserve the integrity of total RNA.

### Single-cell RNA-seq assays

The workflow of these assays is outlined in Figure 1. The samples were used for a first scRNA-seq experiment (SC1) using the Chromium Next GEM Single Cell 3ʹ Reagent Kits v3.1 (Dual Index) protocol with Feature Barcode technology for Cell Multiplexing (10x Genomics). We created two pools (CTRL and HD) from samples; briefly, one of the aliquots of each cryopreserved sample was thawed and washed following the Demonstrated Protocol CG00039 RevD “Fresh Frozen Human PBMC for scRNA-seq” (10xGenomics), and subsequently sorted in a FACSAria III (Becton Dickinson) to remove cell debris and platelets based on size and scatter properties (Supplementary Fig.1) and to collect equal number of cellular events (∼4,500). Next, CTRL and HD pools were processed with the Chromium Next GEM Single Cell 3ʹ Reagent Kits v3.1 (Dual Index) protocol to obtain single cell barcodes for sequencing. Then, we built the DNA libraries and assessed their quality according to the aforementioned protocol. Another experiment was conducted with the same samples (SC2) using the same protocol with the exception of including a prior labelling procedure for each individual sample using Cell Multiplexing Oligo (CMO) tags linked to cell-associated barcodes. The resulting libraries were sequenced in Illumina NovaSeq 6000 system at the Genomics Unit of the Cabimer to achieve 25,000 reads/cell (3’-GEX libraries) and 5,000 reads/cell (CMO libraries).

**Figure 1.**
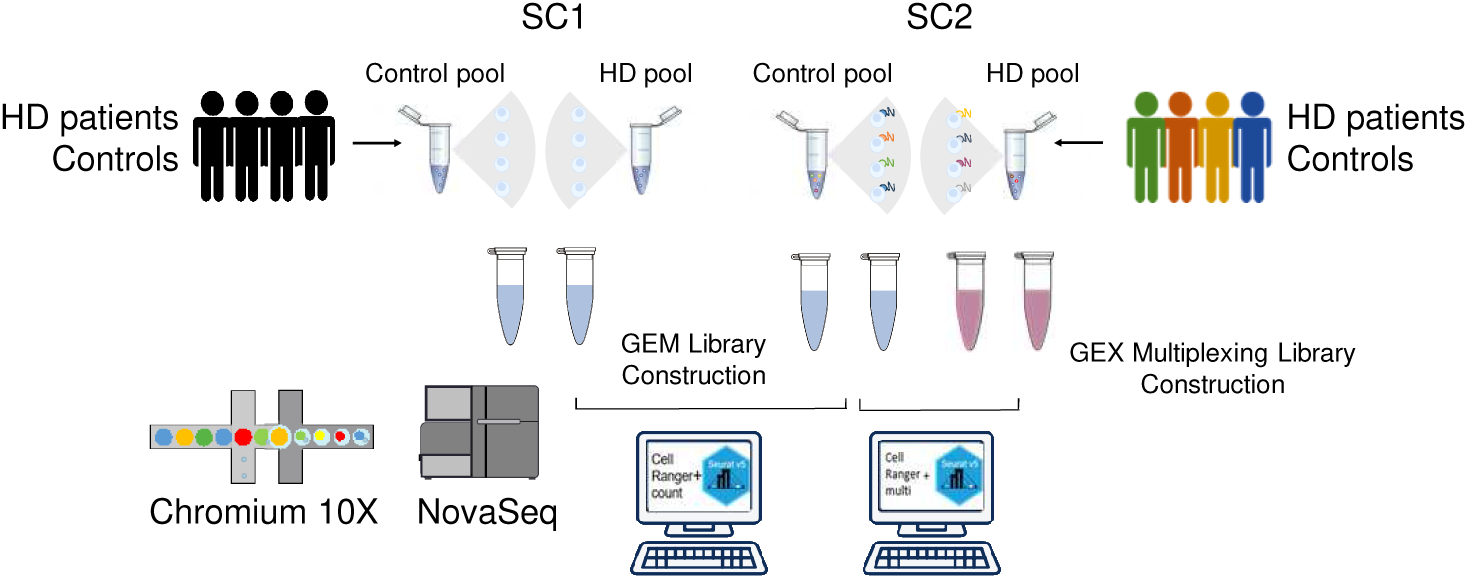
Workflow of the single-cell RNA-seq assays.

### Bioinformatics analysis and statistical analysis

For alignment with the human reference genome GRCh38, we used the 10x Genomics Cell Ranger (v7.2) software. Whereas the ‘cellranger count’ pipeline was used for both experiments, the ‘cellranger multi’ pipeline was also used in SC2 to track single transcriptomes from each individual, adjusting the ‘min-assignment-confidence’ parameter to 0.7 to assign more cells to the tags.

Subsequent analyses were conducted using the “Seurat” package in R (v4.0.4) [20]. The single-cell transcriptomes were filtered according to the following quality control considered metrics: number of unique genes (UMIs) detected per cell (between 800 and 6,000), total number of molecules (reads) within a cell (between 2,000 and 30,000), and percentage of reads mapped to the mitochondrial genome (<15%). The data were then normalized with the “NormalizeData” function applying log-normalization by default. The “scDblFinder” package was used to identify and remove potential doublets (cells with an estimate doublet density score per cell >3) from the dataset. Following doublet filtering, highly variable features were identified (“FindVariableFeatures”), scaled (“ScaleData”) and subjected to Principal component analysis (PCA) (“RunPCA”) in which a dimensionality reduction and clustering pipeline was then applied using the first 20 Principal Components. Cell–cell neighborhoods were computed using “FindNeighbors” function, and clustering was performed using the Louvain algorithm via “FindClusters” with a resolution parameter set to 0.5.

To explore cellular heterogeneity, we used markers described in “Azimuth” [21] to identify which cluster corresponded to each cell type. After trial and error, we opted for grouping minor cell subpopulations that shared cell-specific markers, leading to these groups: CD14^+^ monocytes (*S100A9, CTSS, S100A8, LYZ, VCAN, S100A12, IL1B, CD14, G0S2, FCN1*), other myeloid cells that included dendritic cells (*CTSS, FCN1, NEAT1, LYZ, PSAP, S100A9, AIF1, MNDA, SERPINA1, TYROBP, CD74, HLA-DPA1, HLA-DPB1, HLA-DQA1, CCDC88A, HLA-DRA, HLA-DMA, CST3, HLA-DQB1, HLA-DRB1*), CD4^+^ T-cells (*IL7R, MAL, LTB, CD4, LDHB, TPT1, TRAC, TMSB10, CD3D, CD3G*), other T-cells (*CD3D, TRDC, GZMK, KLRB1, NKG7, TRGC2, CST7, LYAR, KLRG1, GZMA*), NK-cells (*NKG7, KLRD1, TYROBP, GNLY, FCER1G, PRF1, CD247, KLRF1, CST7, GZMB*), naïve B-cells (*IL4R, TCL1A, YBX3*) and active B-cells that included intermediate and memory cells (*COCH, SSPN, LINC01781, NEK6, LINC01857*). Such grouping also enhanced the statistical power of the differential expression analysis within each cell subpopulation (see below). As additional quality controls of our datasets, we also examined the distribution of markers related to plasmablasts (*IGHA2, MZB1, TNFRSF17, DERL3, TXNDC5, TNFRSF13B, POU2AF1, CPNE5, NT5DC2*) as too many short-lived plasmablasts might indicate unrelated infections or vaccinations challenges that might distort the interpretation of the analysis, and platelets (*PPBP, PF4, NRGN, GNG11, CAVIN2, TUBB1, CLU, HIST1H2AC, RGS18, GP9*) to assess the efficiency of our cell sorting procedure. Gene expression patterns of independent cell-specific markers were visualized with the “DotPlot” function from “Seurat” (v5.3.0), splitting by the genotype variable.

Trajectory inference and pseudotime analyses were performed in R (v4.5.0). Cell trajectories were reconstructed with “slingshot” (v2.16.0) on defined clusters [22]. UMAP embeddings were obtained from the “reducedDims” slot, plotted using the color scale of “viridis” (v0.6.5) to represent pseudotime, and overlaid the inferred lineages. Gene dynamics along pseudotime were modelled using “tradeSeq” (v1.22.0) with generalized additive models (GAMs) [23]. The number of knots was determined automatically and consistently set to 10 across all the clusters. Markers of immune cell differentiation stages were found among the genes defining the trajectories (retrieved with “associationTest” function with Benjamini–Hochberg correction).

Within each cellular subpopulation, we performed the differential expression analysis between CTRL and HD using the “FindMarkers” function of the “MAST” package (https://github.com/RGLab/MAST/). Due to different representation of donors despite pooling the same number of cells for each individual for the oil encapsulation process, we considered sex as a latent variable to minimize the retrieval of genes from sex chromosomes (see Results for the uneven distribution of the female marker *XIST*). We considered as differentially expressed genes (DEGs) between controls and patients in each defined subpopilation when the adjusted p-value ≤ 0.05 (using the Benjamini-Hochber method by default). With these results, we considered those DEGs as “Specific” for a particular cellular subpopulation if they were not found in any other pair-wise comparison between control *vs*. patient in the rest of subpopulations (detected in Venn diagrams (https://bioinfogp.cnb.csic.es/tools/venny/, https://bioinformatics.psb.ugent.be/webtools/Venn/); on the contrary, they were considered as “Shared” if found in at least two pair-wise comparisons.

DEGs were compared with those already reported for bulk peripheral blood and brain as they appeared in the Supplementary Tables of the corresponding publications (see Results for further details).To determine whether our DEGs were expressed in human brain, we used the external scRNA-seq datasets from GSE152058 consisted of human caudate nuclei [13]. First, these datasets were processed using the same pipeline applied to our in-house data, using the cellular clusters proposed in the original publication. The “FindAllMarkers” function identified those genes more overexpressed in each cluster (log_2_ fold change > 1, adjusted p-value < 10^-99^). Then, we compared the resulting lists of genes with the whole list of DEGs from PBMCs.

Additional statistical analyses (Mann Whitney U-tests, Spearman’s rank-order correlation coefficients) were conducted in native R environment (v4.4.0).

### Behavioural and gene expression analysis of R6/1 strain

We previously showed in the transgenic B6.Cg-Tg(HDexon1)61Gpb/J strain (*aka* R6/1) that transcriptional dysregulation of HD-related genes showed tissue-specific associations with certain pathological phenotypical traits (see [24] for further details). For the current study, we analyzed the rotarod and weight measurements made in 11 and 13 weeks-old animals and in the striatal and blood samples extracted five days after the behavioural battery (14 weeks-old) in our previous study [24]. Quantitative PCR was performed in QuantStudio 12 K Flex (Thermo Fisher, Madrid, Spain)using HOT FIREpol EvaGreen qPCR Mix Plus 5X (ROX) (Solis Byodine, Tartu, Estonia). The PCR cycling conditions were as follows: 95 °C for 15 min and 40 cycles of 95 °C for 15 s, 60 °C for 20 s, 72 °C for 20 s. Each independent reaction was normalized to the level of *Eef2* [25], and the relative quantitative values were calculated according to the 2^−ΔΔCT^ method. The sequences of all primer pairs are: *Dse*, 5’-GCCTTACTCACTGGAAGTCT-3’ and 5’-TGCTTAGAACTTGCTTGGTC-3’; *Ifit2*, 5’-TTTGACACAGCAGACAGTTA-3’ and 5’-TCCAGTGACTCCTTACTCGT-3’; *Ifitm1*, 5’-CTGAGATCTCCACGCCTGAC-3’ and 5’-CCAGTCGTATCACCCACCAT-3’; *Irf7*, 5’-AAGCATTTCGGTCGTAGGGAT-3’ and 5’-AGTTCGTACACCTTATGCGGA-3’; *Mx1*, 5’-GGTCCAAACTGCCTTCGTAAA-3’ and 5’-CGGATCAGGTTTTCAGCTTCC-3’; *Rsad2*, 5’-AGGACAGGGGTGAATACTTGG-3’ and 5’-CACGGCCAATCAGAGCATTAA-3’; *Samsn1*, 5’-AGGAGAAACACCAAAAACCG-3’ and 5’-ATTCCGAAAACGGTCAAAAT-3’.

## Results and discussion

### scRNA-seq technology detected mild changes in peripheral blood of symptomatic patients that were partially confirmed in published bulk transcriptomics studies

To determine the most altered PBMC subpopulations in HD patients, we performed a first scRNA-seq assay (SC1) comparing two pools of cells from symptomatic donors and controls without the CAG expansion, balanced in sex and age (Fig. 1). After assessing quality (Supplementary Fig. 1), we retained 7761 cells from CTRL and 6788 cells from HD pool, which contained a mean ± SD of transcripts per cell of 9019.716 ± 5103.898 and 8516.413 ± 4922.634, respectively. Despite pooling the same number of cells from each donor (two women and two men), the results of a preliminary differential expression analysis were jeopardized by genes belonging to chromosomes X (e.g., *XIST*, *RPS4X*, *RPL36A*, *TMSB4X*) and Y (e.g., *RPS4Y1*, *USP9Y*, *DDX3Y*, *UTY*, *EIF1AY*, *PRKY*) (see the example of *XIST* expression across clusters in Fig. 2A). These observation indicated that upstream processing and analysis (e.g., storage and sample preparation, cellular encapsulation and labelling, quality filtering of the transcriptomes) might produce stochastic effects possibly amplified by the technical noise that is characteristic of low starting material in single-cell transcriptomics [26], leading to differential contributions (and distributions) of cells from each donor across the resulting clusters. To confirm this, we performed a second scRNA-seq assay (SC2) using the same samples and same batch of reagents, with the only difference of tagging the individual samples using CMOs prior to pooling for single cell labelling (Fig. 1). As observed in Fig. 2B, the distribution of CMOs was also uneven across clusters in both CTRL and HD pools. For this reason, we considered sex as an additional variable to minimize the influence of sexual chromosomes in subsequent analyses.

**Figure 2.**
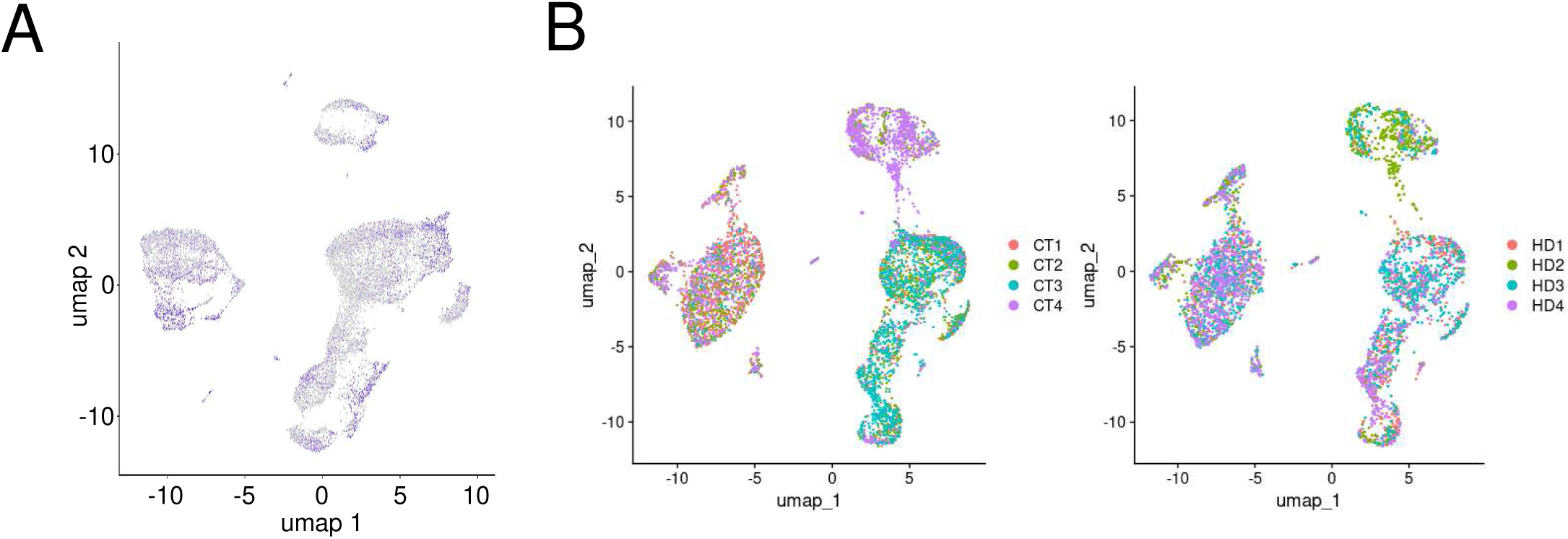
Different contribution of donors to cellular clusters. *A*, UMAP with the distribution of *XIST* expression (SC1). *B*, UMAP with the distribution of cells per donor, as tagged using CMOs prior to sample pooling (SC2).

Taking profit of the high quality of SC1 in which filtered cells from CTRL and HD pools showed highly similar distribution of UMIs and total reads per cell (Supplementary Fig. 1), we clustered the individual transcriptomes of this assay into relevant cell subpopulations (T-cells, NK-cells, myeloid cells, and B-cells) grouping cellular subtypes with low abundance of cells according to the distribution of common markers in a balance to reduce redundancy while preserving cellular identity (Fig. 3A); in this way, we reduced the complexity of the data while increasing the statistical power of downstream differential expression analysis. We did not observed substantial differences in the proportion of cells across clusters between both genotypes (Fig. 3B). The resulting proportions (75.5 % of lymphocytes consisted of 82.7% of T cells, 10.7% of B cells, 6.6% of NK cells, and 24% of myeloid cells including monocytes and dendritic cells) fell within the expected range [27] meanwhile a very small fraction of unwanted cell types (plasmablasts and platelets, 0.4% and 0.1%, respectively) was obtained. We confirmed the annotation of the resulting cell subpopulations by plotting specific markers for T-cells (*CD3E*, *CD28*), B-cells (*CD19*, *MS4A1*), NK cells (*NCAM1*, *KLRC1*) and macrophages/monocytes (*TNFSF13*, *LILRB3*), not used for cluster identification (Fig. 3C). Based on the UMAP, we performed a trajectory analysis in the three major clusters of cells (all T and NK cells, all myeloid cells and all B-cells) with sufficient number of cells to reflect the dynamics of maturation and differentiation of peripheral immune subpopulations in the SC1 experiment: as observed in Fig. 3D, the distribution of cells across the global trajectories was highly similar between genotypes.

**Figure 3.**
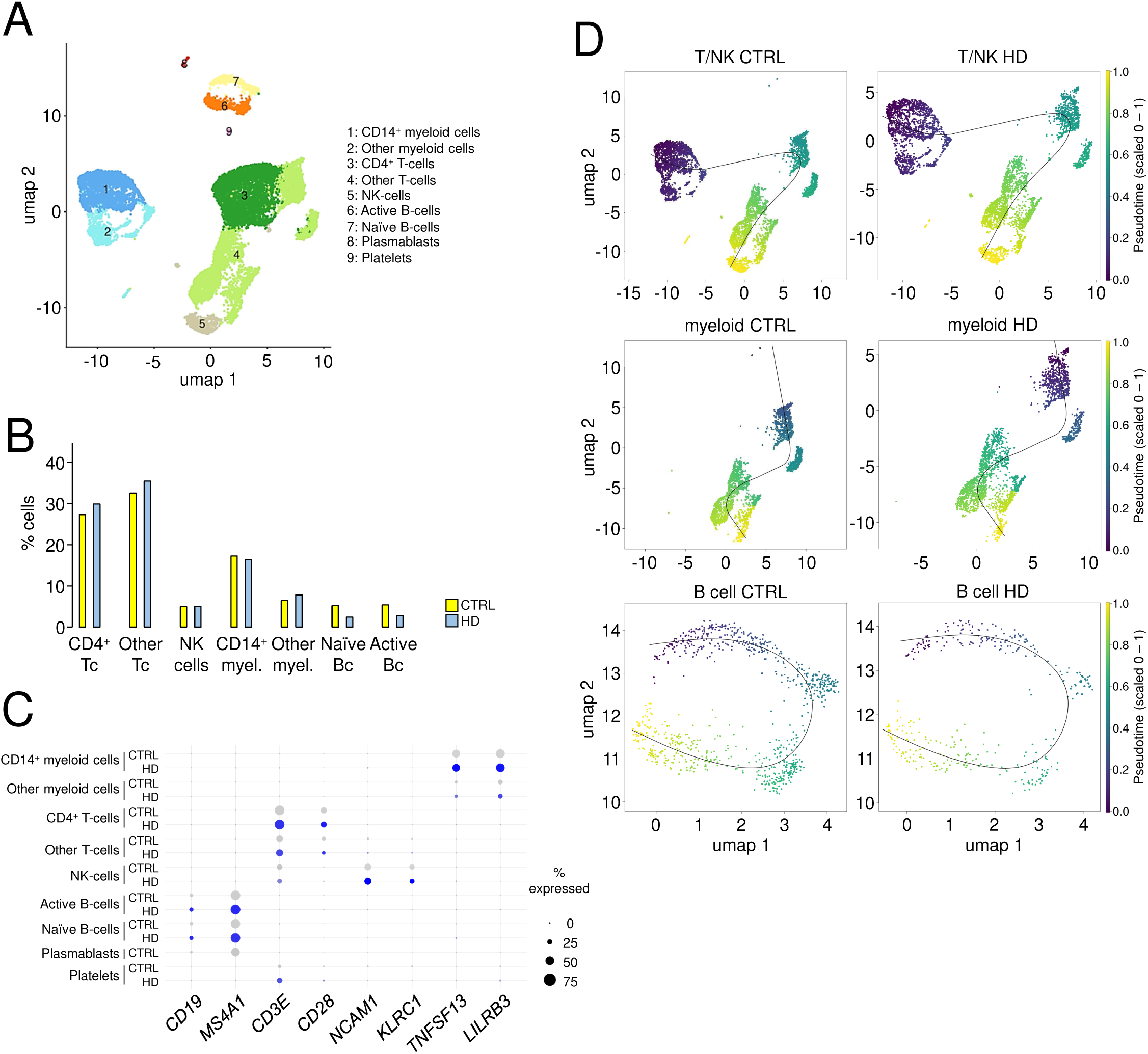
General results of single-cell RNA-seq assay in PBMCs from HD patients and control donors. *A*, UMAPs showing the PBMC subpopulations from SC1. *B*, Percentage of cells for each subpopulation from the total that passed the quality filters for each pool of controls (CTRL) and patients (HD). *C*, Dot plot of cell-specific markers across the defined cellular clusters in both pools. *D*, UMAPs of CTRL and HD cells across the global trajectories defined in major cell clusters, scaled from 0 to 1.

Next, we conducted the differential expression analysis between control *vs*. patient cells in each cell subpopulation depicted in Fig. 3A. Although important fractions of DEGs in patients related to controls were apparently specific of the defined cell subpopulations (i.e., only appearing in a single pair-wise comparison between controls and patients), ∼52% of upregulated genes and ∼35% of downregulated genes were detected in more than one population (Fig. 4A). Among the most frequent DEGs across cell subpopulations (appearing at least in five pair-wise comparisons between controls and patients), we found the following genes: the eukaryotic translation initiation factor kinase *EIF2AK2*, the endoplasmic reticulum aminopeptidase *ERAP2*, the epithelial stromal interactor *EPSTI1*, the interferon-induced *IFI44*, *IFI44L* and *IFIT3*, the ubiquitin-like modifier *ISG15*, the mitochondrial-like *MTRNR2L12,* the dynamin GTPases *MX1* and *MX2*, and the interferon-induced apoptotic *XAF1* as upregulated genes, together with the haemoglobin *HBB*, the ribosomal *RPS26* and the adhesion G-protein coupled receptor *ADGRG7* as downregulated genes in patients. No information has been found regarding these genes in HD, although genetic variants of *EIFAK2* have been linked to dystonia [28].

**Figure 4.**
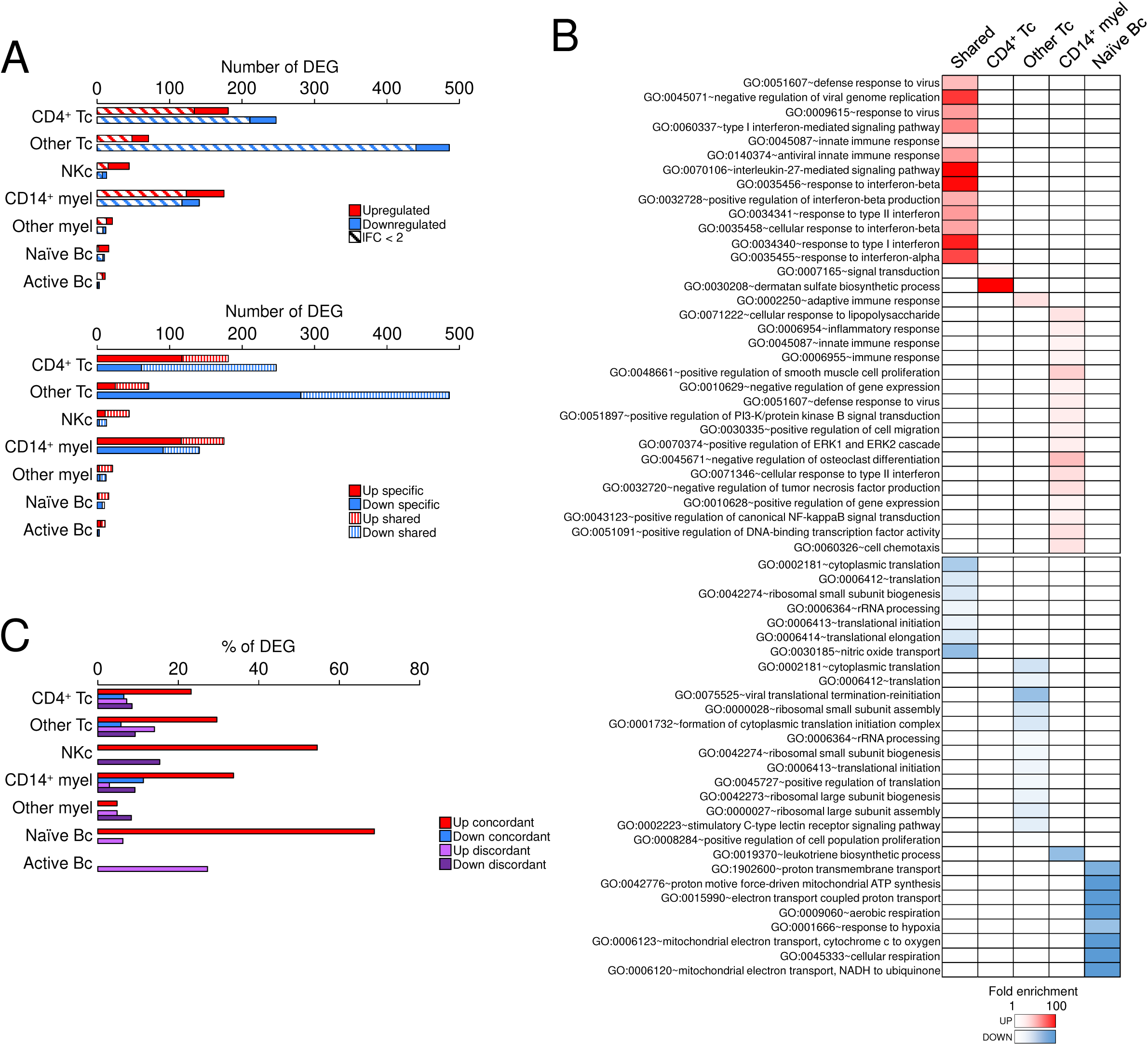
Differential expression analysis in PBMC subpopulations between HD patients and control donors. *A*, Number of differentially expressed genes (DEGs) between CTRL and HD in each subpopulation and assay (adjusted p-value < 0.05, MAST). Upregulated and downregulated genes in patients’ cells compared to controls’ cells are shown; “Specific” refers to significant genes that are only found in this subpopulation whereas “Shared” refers to genes observed in at least two subpopulations. *B*, Significant Gene Ontology (GO) enriched terms (FDR < 0.05) obtained from specific and shared DEGs for each direction of change (UP and DOWN). No significant term was retrieved in “Other myeloid” and “Active Bc”. *C*, Percentage of DEG in each cell subpopulation that was reported in at least one publication [5,6,11,38–40,49]. “Concordant” and “discordant”, in the same or opposite direction of change, respectively.

Unsurprisingly, upregulated genes were mostly related with immune response (innate, interferon-mediated, virical), showing minor functional specificity (e.g., chemotaxis in CD14^+^ monocytes) (Fig. 4B). Downregulated genes were mainly enriched in protein translation and ribosomal biogenesis, which is intriguing considering that these processes are highly energy-demanding processes that require fine-tuning regulation under T cell activation and subsequent proliferation [29–31], and their components are upregulated in B-cell maturation, probably to face antibody production [32]. Other enriched functions were mitochondrial genes in naïve B-cells and leukotriene metabolism in CD14^+^ monocytes; the latter pathway produces important mediators of inflammatory and allergic responses that can be therapeutically targeted to modulate neuroinflammation and oxidative stress in neurodegenerative conditions including striatal neurotoxicity [33–36]. Whether the downregulation of components of these processes in HD PBMCs was part of a homeostatic response to limit the hyper-reactivity reported in peripheral immune cells from HD patients and mouse models [7,8,11,12,37] deserves further attention.

To confirm the potential alterations observed in the scRNA-seq assays, we examined whether some of the retrieved DEGs were also reported in previous studies using larger cohorts. Since several changes were retrieved across multiple cell types (Fig. 4A), we reasoned that they should be detected in bulk transcriptomics studies. Indeed, the percentage of DEGs in the scRNA-seq that were also found in any of these studies varied an average of ∼13% (Fig. 4C). More precisely, overlapping upregulated genes were more frequent and concordant than overlapping downregulated genes (Fig. 4C and Table 1), being the most frequent genes the interleukin *IL1B* and the phospholipid scramblase *PLSCR1* appearing in three of the reported studies. Of the changing genes, ∼22% of the upregulated genes were not cell-specific according to our cut-offs, which justified their detection in whole blood experiments.

**Table 1.**
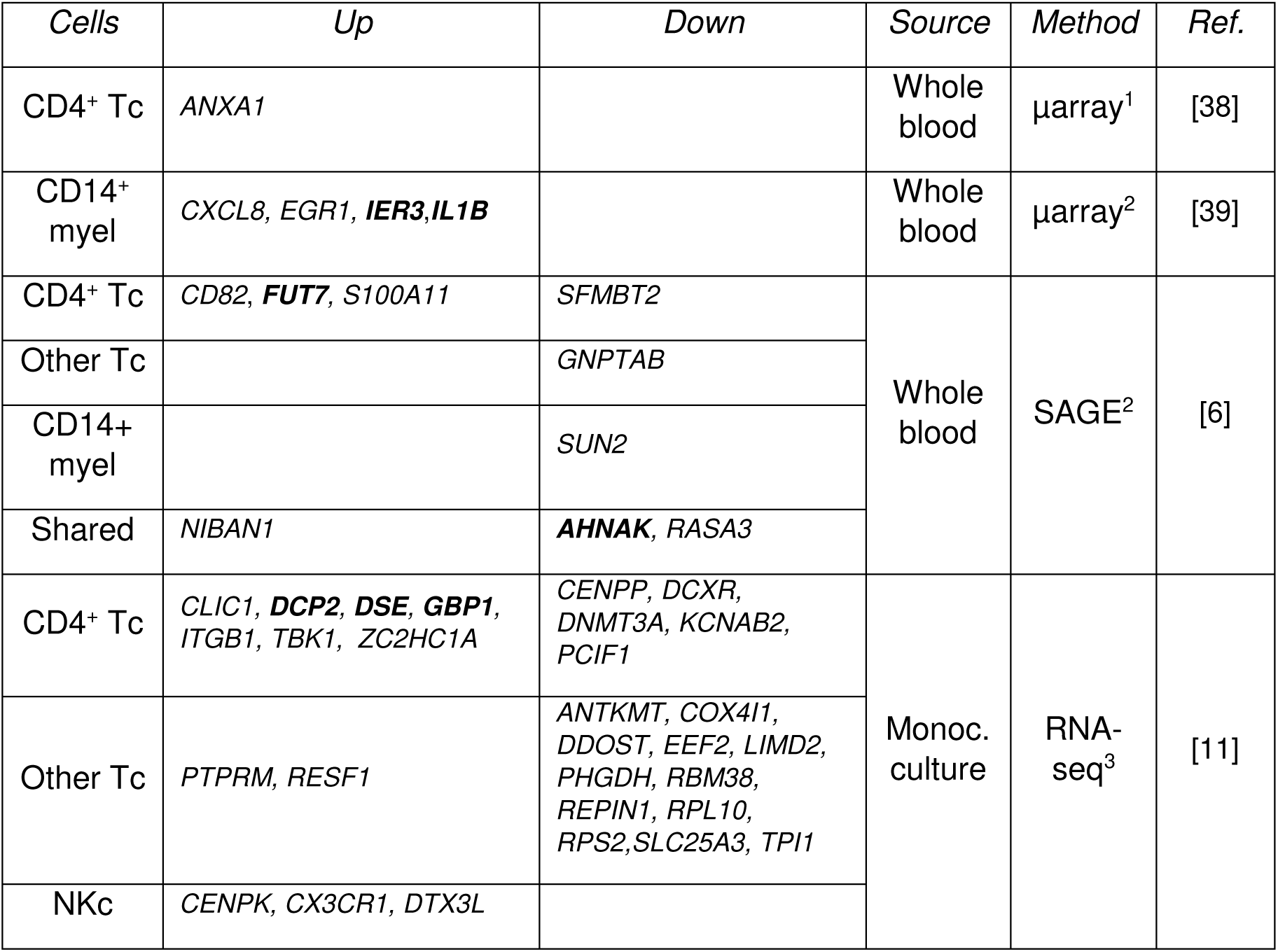
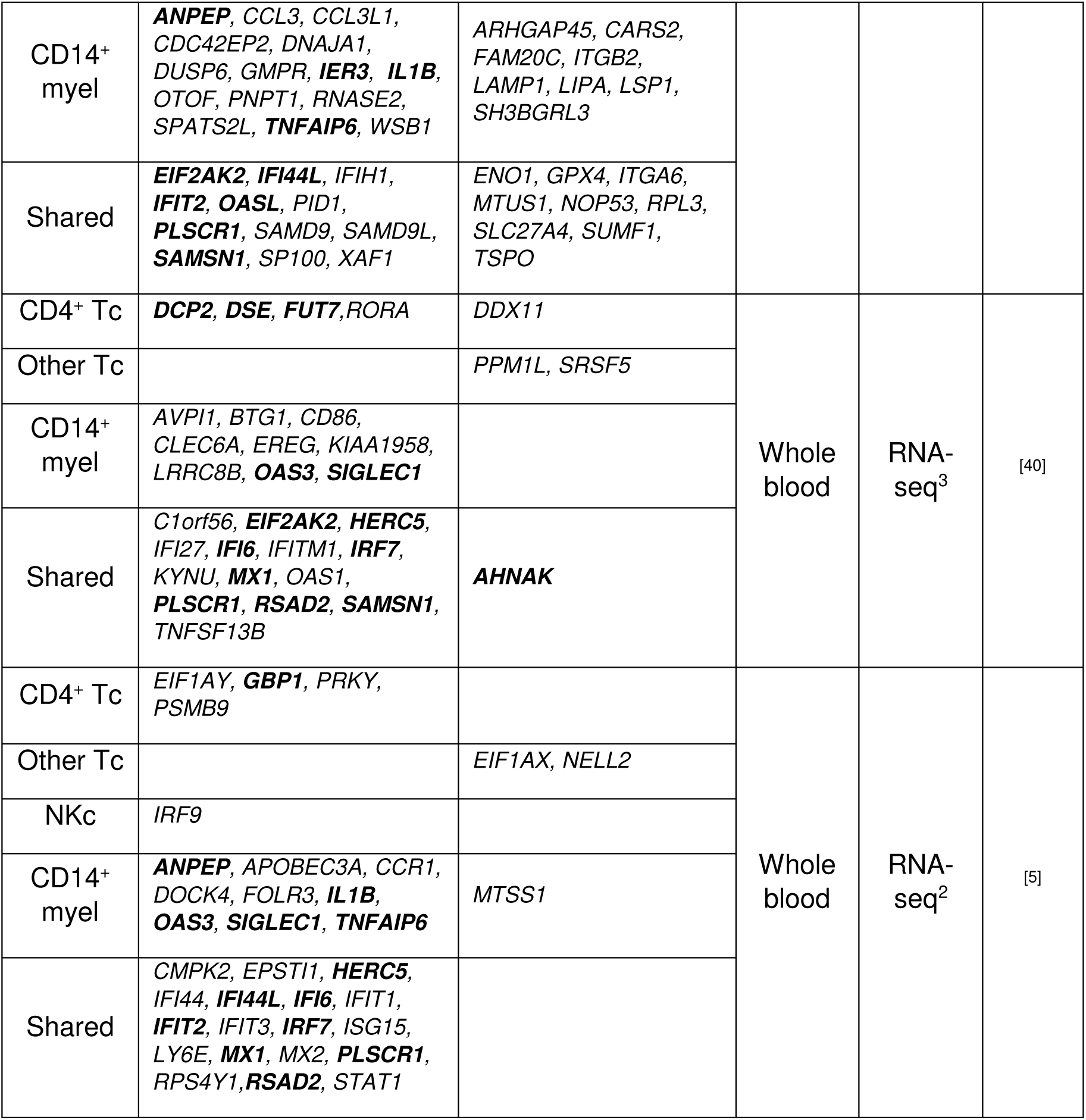
Common differentially expressed genes (DEGs) between scRNA-seq results of this study and those published from bulk transcriptomics. ^1^unadjusted p-value < 0.0005; ^2^adjusted *p*-value < 0.05; ^3^unadjusted *p*-value <0.05; Bold, common DEGs in at least two published datasets in any cell subtype. SAGE, Serial Analysis of Gene Expression.

### Using a controlled and phenotypically characterized mouse model for potential validation of novel biomarkers

Next, we asked whether the described peripheral blood alterations in HD patients can be linked to disease progression as potential biomarkers. To this aim, we took profit of a previous phenotypical characterization of early symptomatic R6/1 mice and accompanying samples from the same animals; briefly, mice were subjected to a battery of behavioural tests that also included measurements of certain phenotypical traits at two time points (11 and 13 weeks-old) to assess progression. Next, samples were collected five days after the last measurements (to avoid the potential influence of genes related with handling stress) to examine brain changes in gene expression that could be correlated with the manifested pathological traits across individuals; for example, the altered expression patterns of striatal *Penk*, *Plk5* and *Itpka* genes were mildly correlated with poor performance in the accelerated rotarod and weight loss [24]. Together with the dissection of brain tissues, we also collected blood samples of the same reported animals to validate whether the gene expression patterns observed in total blood and PBMCs of patients might be correlated with the phenotypical traits measured in mouse. For these assays, we selected genes that were found to be expressed in the most abundant PBMC cell types (Table 1). Of these, only *Ifit2* exhibited a trend towards upregulation, whereas other genes were downregulated in R6/1 samples (most prominently *Rsad2*), indicating that there were species-specific differences at the peripheral level (Fig. 5A). Alternatively, transcriptional changes in human blood were too mild, as evidenced by the extensive use of unadjusted *p*-values in global-wide studies [40–42], to be well reproduced in mouse models.

**Figure 5.**
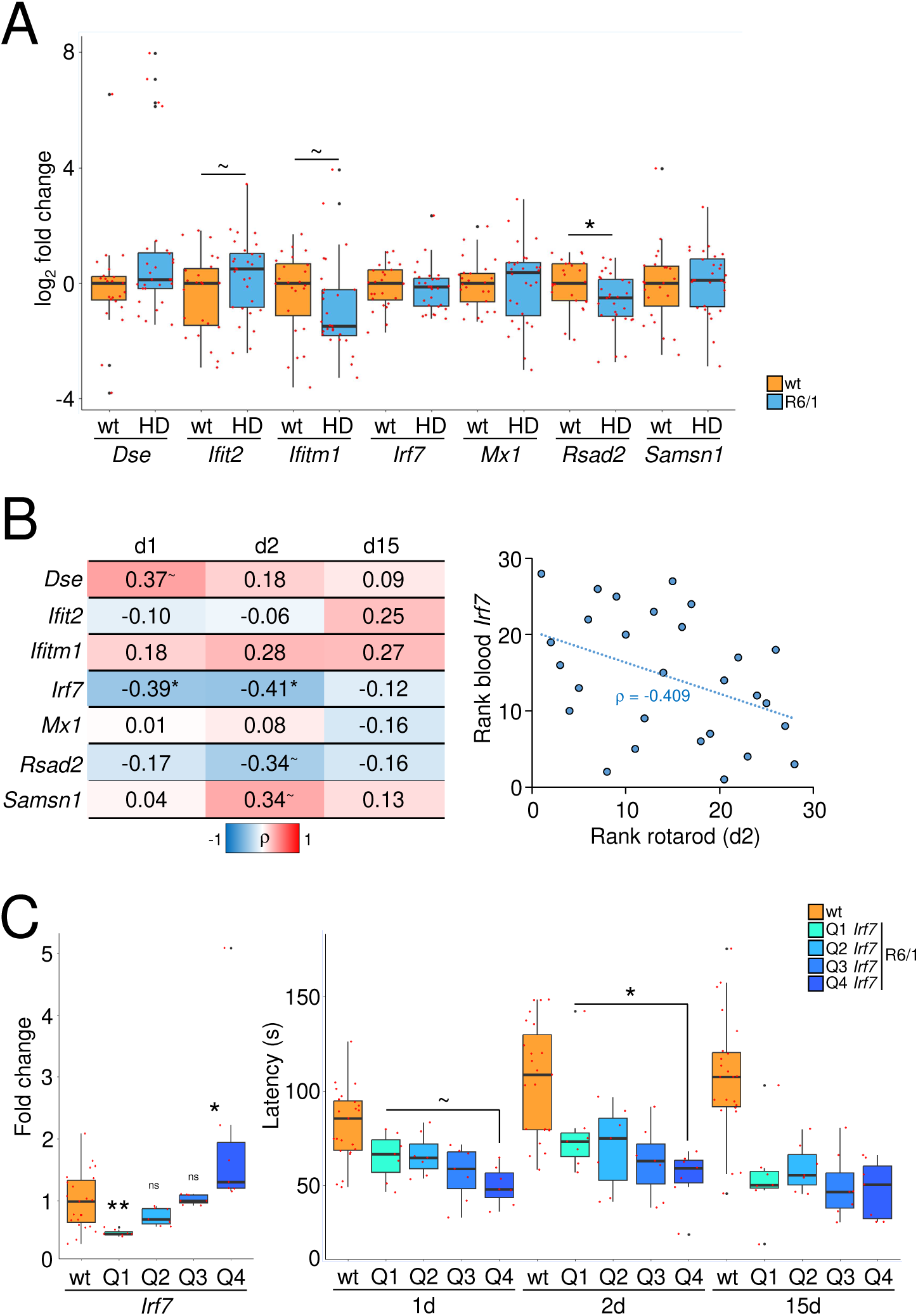
Correlation between blood gene expression and motor performance in the R6/1 mouse model. *A*, Box-and whisker plots showing the fold changes in mutant mice (HD, n = 28) compared to wild-type littermates (wt, n = 23) for the indicated genes in blood. *, *p*-value < 0.05; ∼, *p*-value < 0.1 (Mann-Whitney U-test). *B,* Left, heatmap of Spearman correlation coefficients (ρ) between the examined transcripts in blood from R6/1 mice and rotarod performance at day 1 (d1), 2 (d2) and 15 (d15) (see [24]) (*, *p*-value < 0.05; ∼, *p*-value < 0.1). Right, correlation between peripheral *Irf7* expression and latency on the rotarod. *C*, Box-and whisker plots showing the rotarod performance (measured as latency in s) for the groups of animals (right) according to the expression changes of *Irf7* in blood (left). ns, not significant; *, *p*-value < 0.05; **, *p*-value < 0005, Mann-Whitney U-test referred to wt (left) and between quartiles Q1 and Q4 (right).

In any case, gene expression might still exhibit relevant differences across individuals that might be reflected at the phenotypical level. Focusing on the rotarod performance, the most well-known paradigm to assess impaired motor coordination in HD mouse models, we observed that motor impairment was significantly worsened as *Irf7* became upregulated in mutant mice (Fig. 5B-C). IRF7 is member of the interferon regulatory transcription factors that recognizes the interferon (IFN)-stimulated response element (ISRE) in the DNA to mediate the response of IFN type I and III in viral infections but also in autoimmune diseases and cancer [43,44] but its precise role in HD has not investigated yet. This correlation was significant when comparing the rotarod latencies at an early stage (11 weeks old) which was later lost (13 weeks old) due to a rapid decline in this phenotypical trait (see [24] for further details), leveraging this behaviour in all mutant mice independently on the *Irf7* alteration (Fig. 5B-C) and other striatal genes [24]. No correlation was observed with weight, which loss is also a hallmark of HD (Supplementary Fig. 2). Overall, these results suggested that only few examples of peripheral transcriptional changes were susceptible to be linked to measurable manifestations of the disease.

### Blood alterations in gene expression can be found in the brain

To establish a potential link between brain events and altered patterns of peripheral biomarkers, we investigated whether the DEGs defined in the scRNA-seq approach were found in published datasets obtained from post-mortem caudate nuclei [18] (unadjusted *p*-value < 0.001) and prefrontal cortex [45] (adjusted *p*-value < 0.05), revealing that ∼38% of upregulated genes and ∼20% of downregulated genes in patients’ PBMCs were also disrupted in the same direction of change in patients’ postmortem brain (Fig. 6A). This was plausible because these peripheral DEGs were mainly expressed in non-neuronal cells, mainly in brain resident T cells, microglia, and vascular cells, according to published scRNA-seq data from human caudate nuclei [13] (Fig. 6B).

**Figure 6.**
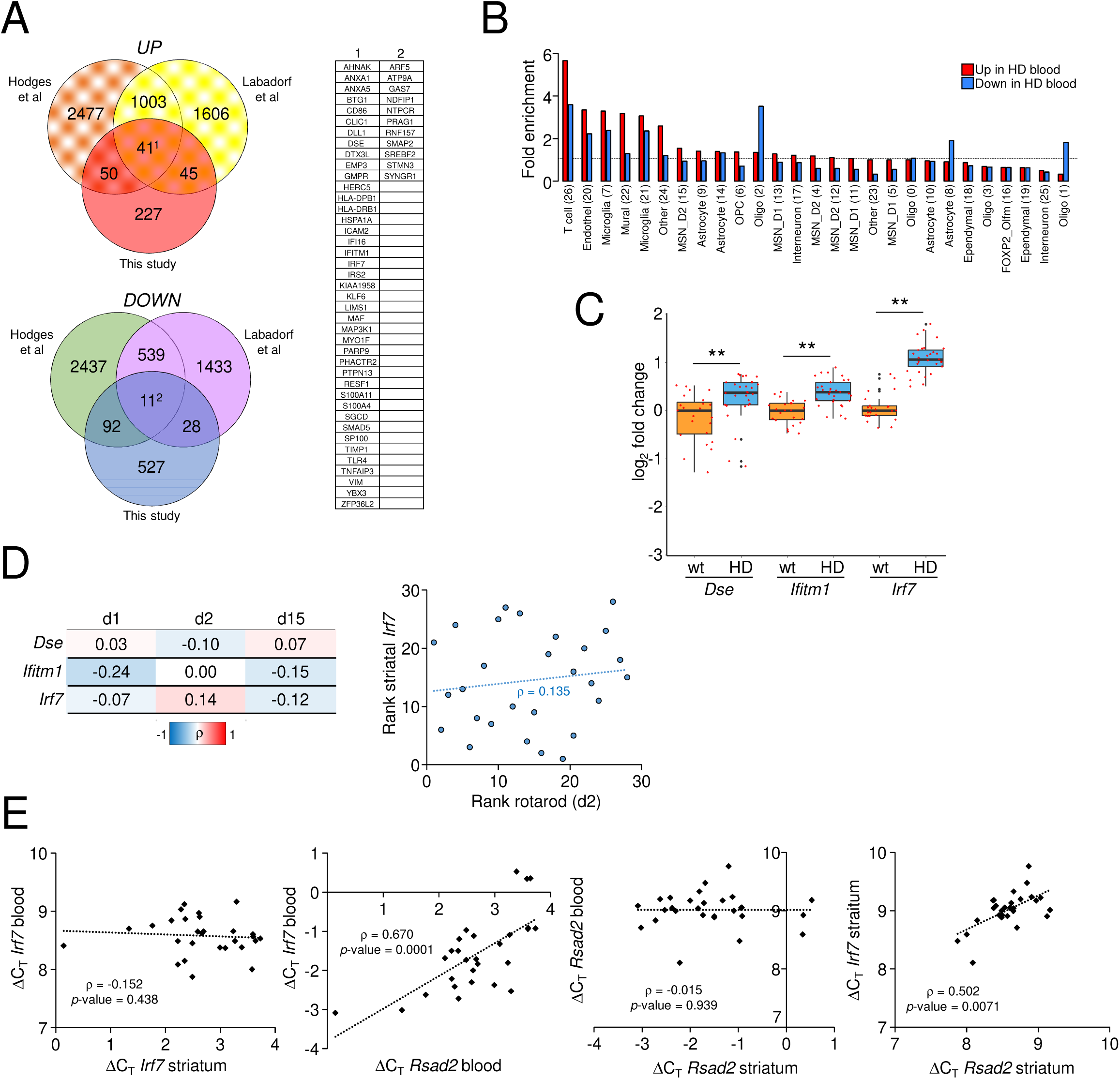
Analysis of blood transcriptional counterparts in the brain of patients and R6/1 mice. *A*, Venn diagram with the overlapping DEGs between scRNA-seq data from patients’ PBMCs (“This study”) and published bulk RNA-seq data from post-mortem patients’ brains (“Hodges et al” [18] and “Labadorf et al” [45]). Besides, table showing the overlapping genes in all studies following the numbering of the Venn diagram. *B*, Fold enrichment analysis of expression of peripheral DEGs across cell subtypes as defined in [13] for human caudate nucleus (number of cluster between parenthesis). *C*, Box-and whisker plots showing the fold changes in mutant mice (HD, n = 28) compared to wild-type littermates (wt, n = 23) for the indicated genes in striatum. **, *p*-value < 0.005, Mann-Whitney U-test. *D*, Left, heatmap of Spearman correlation coefficients (ρ) between the examined transcripts in striatum from R6/1 mice and rotarod performance at day 1 (d1), 2 (d2) and 15 (d15) [18]. Right, correlation between striatal *Irf7* expression and latency on the rotarod. *E*, Gene expression is represented as ΔC_T_ (C_T_ gene – C_T_ *Eef2*) and correlation as Spearman correlation coefficient (ρ) and associated *p*-value.

Taking profit of available striatal samples from the same R6/1 mice [24] from which blood was examined (Fig. 5), we confirmed the upregulation of *Dse*, *Ifitm1* and *Irf7* in the mouse model that was much more consistent compared to blood, being *Irf7* the most changing gene (Fig. 6C). Next, we evaluated whether the striatal expression of these genes were correlated with both peripheral levels and phenotype. However, we did not observe any correlation with motor impairment, either for *Irf7* (Fig. 6D). Notably, the expression of *Irf7* and *Rsad2*, which was also expressed in the human and mouse brains, were mutually correlated within each compartment (either striatum or blood) but the expression of each gene was not correlated across compartments (Fig. 6E), strengthening the view that brain and peripheral alterations of the same genes were largely dissociated as they are likely to be expressed under the control of distinct regulatory mechanisms, influenced by the tissue context. Taking into account that the transgenic R6/1 strain is an aggressive model which use could be an important limitation of our study, slow progressive knock-in-based strains that reproduce both the mHTT production and the loss of a functional HTT allele can improve brain and peripheral correlations and facilitate the comparisons between patients and mouse models, as the manifested phenotype is more similar to the human pathology. However, both transgenic and knock-in HD mouse models do not develop a consistent neuroinflammatory response in the striatum (e.g., gliosis and microglia activation) [13,24,25] in contrast to patients [18,45–47] that may limit the translation of our results from mice to humans. Nonetheless, the mild but significant upregulation of at least three genes involved in immune response in the striatum of a HD mouse model is of interest, which is in agreement with the documented increase of interferon-responsive genes in the striatum of R6/2 and zQ175DN mouse models [13]. These observations may pinpoint to more subtle but still undisclosed roles for these genes in the dysfunction of the basal ganglia in these animals that may have a more general impact on the functioning of the corticostriatal circuitry. For example, complement proteins secreted from locally activated microglia can provoke synaptic loss in presymptomatic stages that can lead to cognitive impairment [48].

### Conclusions

Our results derived from single-cell transcriptomics complemented previous reports aimed at obtaining transcriptional-based biomarkers from peripheral blood of patients to measure the onset and progression of HD. To solve the absence of a cohort of validation, we used published bulk transcriptomics datasets that might be biased towards the validation of genes that were altered in multiple cell types, at the expense of leaving cell-specific changes largely unexplored in this study. As our pools consisted of samples from few donors, we attempted to compensate the lack of correlative evidence based on clinical data by using our published phenotypical data from mouse models that were inbred under controlled genetic and environmental conditions, thus minimizing confounding factors. Despite species-specific divergences, we were able to find moderate correlations with the rotarod performance and (at least) the lymphocytic expression of *Irf7* gene, which striatal counterpart was prominently upregulated, in the mutant mice. However, we found that *Irf7* expression was dissociated between striatum and blood of the same animals, undermining the use of peripheral markers to effectively infer the disease stage in the brain. More exhaustive screens in phenotypically characterized mice may reveal more and better correlated genes, as a first line to propose potential candidates for their further assessment in humans.

## Acknowledgements

We want to acknowledge the patients, the Asociación Valenciana de Enfermedad de Huntington and the Biobank of ISABIAL (adhered to the Spanish National Biobanks Network and integrated in the Valencian Biobanking Network) for their collaboration. We also thank Yolanda Guillén-Montalbán (Universitat Oberta de Catalunya) for their bioinformatics assistance, and Javier Caler and José Mulet (Genomics Unit of Instituto de Neurociencias, CSIC-UMH) for their technical assistance using the Chromium single-cell technology.

## Author contributions

Conceptualization, Funding acquisition, Project administration, Resources and Supervision: L.M.V. Data curation: P.M.C. Formal analysis and Software: P.M.C. and S.O.M. Investigation: P.M.C., J.F.G.S., S.O.M., S.M.M. and L.M.V. Methodology: P.M.C., S.O.M. and L.M.V. Validation: J.F.G.S. and P.M.C. Visualization: P.M.C, S.O.M. and L.M.V. Writing—original draft: L.M.V. Writing—review & editing: L.M.V. and J.F.G.S.

## Data availability

The scRNA-seq data can be downloaded from ArrayExpress using the accession number E-MTAB-15129.

