## Supplementary Figures for "Single-cell transcriptomics and mouse model phenotyping for biomarker screen of peripheral blood biomarkers in Huntington’s disease"

**Supplementary Figure 1. Cell filtering in the scRNA-seq assays.** *A*, The selected window for cell sorting was the same for all samples. *B*, Violin plots showing the parameters of nFeature_RNA (number of unique UMIs / cell), nCount_RNA (total number of reads / cell) and percent.mt (% of mitochondrial reads) for each pool of samples after filtering (see Materials and Methods).

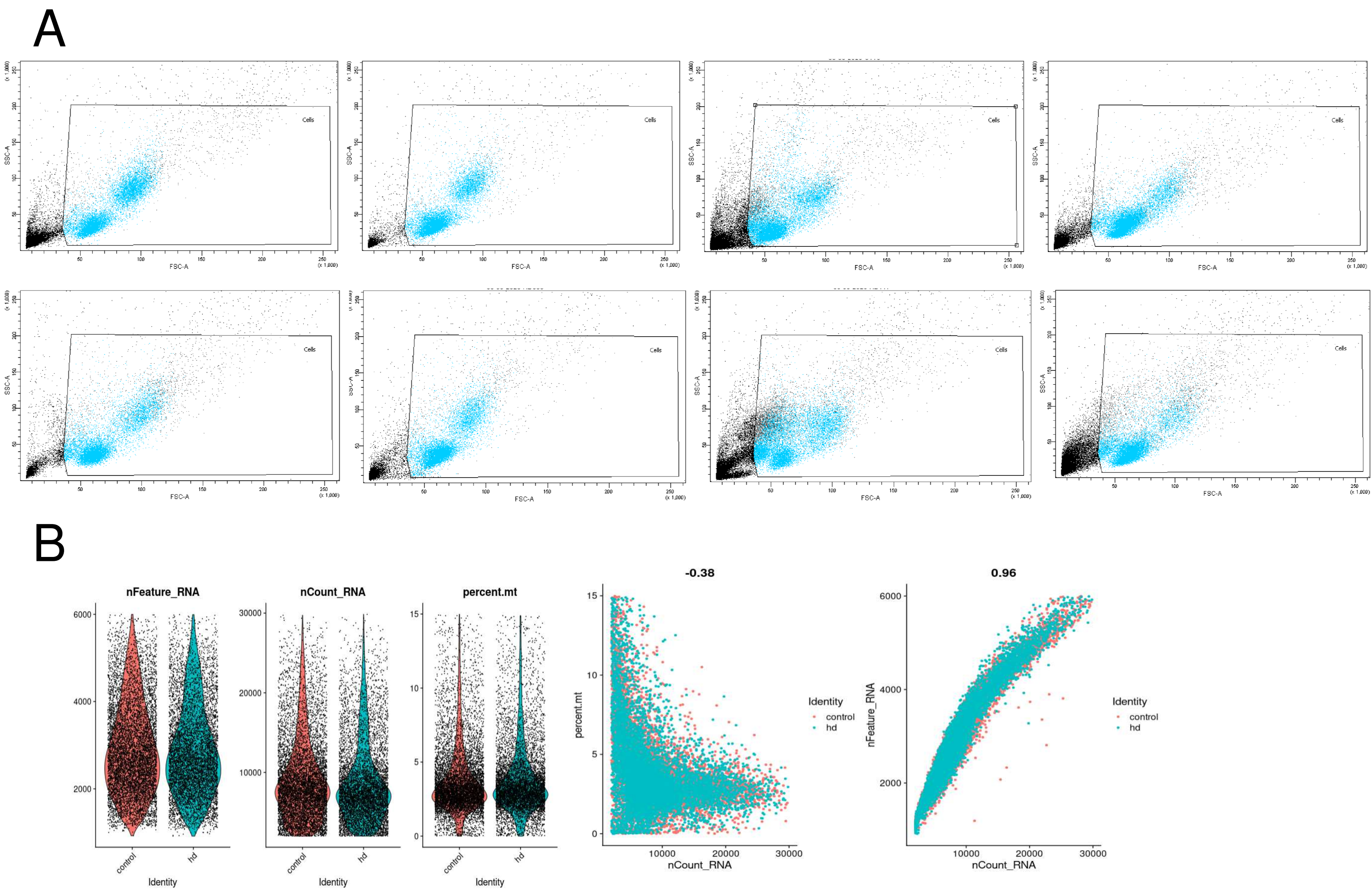


**Supplementary Figure 2. Correlation between gene expression and motor performance in the R6/1 mouse model.** *A*-*B*, Heatmap of Spearman correlation coefficients (ρ) between the examined transcripts in blood (*A*) and striatum (*B*) from R6/1 mice and rotarod performance at 11 (11w) and 13 (13w) weeks.

A

| | 11w | 13w |
|---|---|---|
| *Dse* | 0.17 | 0.05 |
| *Ifit2* | 0.06 | 0.03 |
| *Ifitm1* | 0.20 | 0.08 |
| *Irf7* | -0.11 | -0.01 |
| *Mx1* | 0.27 | 0.30 |
| *Rsad2* | 0.10 | 0.14 |
| *Samsn1* | 0.24 | 0.13 |

-1 ρ 1

B

| | 11w | 13w |
|---|---|---|
| *Dse* | 0.19 | 0.18 |
| *Ifitm1* | -0.16 | -0.18 |
| *Irf7* | 0.04 | -0.03 |

-1 ρ 1
